# Transcriptional regulatory networks that promote and restrict identities and functions of intestinal innate lymphoid cells

**DOI:** 10.1101/465435

**Authors:** Maria Pokrovskii, Jason A. Hall, David E. Ochayon, Ren Yi, Natalia S. Chaimowitz, Harsha Seelamneni, Nicholas Carriero, Aaron Watters, Stephen N. Waggoner, Dan R. Littman, Richard Bonneau, Emily R. Miraldi

## Abstract

Innate lymphoid cells (ILCs) can be subdivided into several distinct cytokine-secreting lineages that promote tissue homeostasis and immune defense but also contribute to inflammatory diseases. Accumulating evidence suggests that ILCs, similarly to other immune populations, are capable of phenotypic and functional plasticity in response to infectious or environmental stimuli. Yet the transcriptional circuits that control ILC identity and function are largely unknown. Here we integrate gene expression and chromatin accessibility data to infer transcriptional regulatory networks within intestinal type 1, 2, and 3 ILCs. We predict the “core” sets of transcription-factor (TF) regulators driving each ILC subset identity, among which only a few TFs were previously known. To assist in the interpretation of these networks, TFs were organized into cooperative clusters, or modules that control gene programs with distinct functions. The ILC network reveals extensive alternative-lineage-gene repression, whose regulation may explain reported plasticity between ILC subsets. We validate new roles for c-MAF and BCL6 as regulators affecting the type 1 and type 3 ILC lineages. Manipulation of TF pathways identified here might provide a novel means to selectively regulate ILC effector functions to alleviate inflammatory disease or enhance host tolerance to pathogenic microbes or noxious stimuli. Our results will enable further exploration of ILC biology, while our network approach will be broadly applicable to identifying key cell state regulators in other *in vivo* cell populations.

## Introduction

Innate lymphoid cells (ILCs) are recently characterized cells that regulate critical aspects of tissue homeostasis, inflammation, and repair (Cherrier et al., 2012; Eberl et al., 2015; van de Pavert et Vivier, 2015; Sonnenberg et Artis, 2015; Spits et Cupedo, 2012; Walker et al., 2013). As the innate counterparts of adaptive T lymphocytes, ILCs lack somatically rearranged antigen receptors and respond rapidly upon encountering specific metabolites or cytokines (Tait Wojno et Artis, 2016; Vivier et al., 2018). In addition to this hallmark feature, ILCs are enriched at mucosal and barrier surfaces where they are poised to coordinate tissue immunity and homeostasis (Tait Wojno et Artis, 2012).

Although recent studies highlight the heterogeneity of intestinal ILCs (Björklund et al., 2016; Gury-BenAri et al., 2016), the cells can nevertheless be broadly classified into three principal groups based on their signature effector cytokines, functional characteristics, and dependence on lineage-determining transcription factor expression (Eberl et al., 2015; Spits et al., 2013; Walker et al., 2013). Type 1 ILCs include ILC1 and natural killer (NK) cells that produce interferon gamma (IFNγ) and require the transcription factor T-bet (*Tbx21*). NK cells also produce IFNγ, but they are unique in their potent cytolytic function and additional dependence on eomesodermin (Eomes) (Pikovskaya et al., 2016). Type 2 ILCs (ILC2s) require GATA3 and secrete the cytokines IL-4, IL-5, and IL-13. ILC3s comprise a heterogeneous group of related cells defined by their dependence on RORgt and production of IL-22 and IL-17 cytokines. During development, the T-helper-like ILCs (ILC1, ILC2, and a subset of ILC3) arise from a common progenitor that diverges from NK cell and lymphoid tissue inducer (LTi) cell progenitors (Constantinides et al., 2014; Klose et al., 2014). Unlike NK cells, these ILCs are dependent on IL7Rα. While some of the TFs driving early lineage commitment of ILCs have been discovered (Constantinides et al., 2014, 2015; Fang et Zhu, 2017; Lim et al., 2017; Serafini et al., 2015; Yu et al., 2013), the transcriptional mechanisms that maintain ILC identity and function are less well studied.

An emerging property of ILCs is their apparent functional and phenotypic flexibility in response to changing tissue environments (Lim et al., 2016, 2017; Melo-Gonzalez et Hepworth, 2017; Ohne et al., 2016). For example, ILC1 and ILC3 exist on a continuum marked at its extremes by the exclusive expression of lineage-specifying factors T-bet or RORγt, while intermediate populations express varying levels of both TFs (Bernink et al., 2015; Klose et al., 2013; Verrier et al., 2016; Vonarbourg et al., 2010). Their interconversion can result in the accumulation of a pro-inflammatory IFNγ-producing “ex-ILC3” that drive intestinal inflammation (Bernink et al., 2013; Vonarbourg et al., 2010). This property mirrors the relationship between Th1 and Th17 cells, with conversion of RORγt^+^ IL-17A-expressing cells into T-bet^+^ IFNγ-producing Th1-like “pathogenic Th17 cells” in the presence of IL-23 (Hirota et al., 2011). Plasticity between ILC1 and ILC2 as well as ILC1 and NK cells has also been reported (Lim et al., 2017). These findings highlight the need for a better understanding of the molecular mechanisms that govern ILC functional plasticity.

Recent studies have described epigenetic and transcriptional signatures for ILC subsets and suggested subset-specific TF regulators based on TF-motif analysis of differentially accessible chromatin regions (Gury-BenAri et al., 2016; Koues et al., 2016; Robinette et al., 2015; Shih et al., 2016). However, the TF regulators of individual genes in each ILC subset remain largely unknown. These regulatory relationships are the building blocks of larger gene programs driving ILC function and require broad, unbiased and systematic investigation.

Transcriptional regulatory networks (TRNs) describe the regulatory interactions between TFs and their target genes. TRN construction is especially challenging in relatively rare *ex vivo* cell types like ILCs, because many transcriptional regulators are unknown, and it is technically difficult to obtain direct TF occupancy data for those that are. In this context, integration of chromatin accessibility with TF motif databases provides a powerful means to identify putative regulatory regions and the candidate TFs binding to those regions (Pique-Regi et al., 2011). We recently developed a version of *Inferelator* algorithm that integrates TF motif analysis of chromatin accessibility and gene expression data to infer TRNs (Miraldi et al., 2018). Importantly, we showed that this method, applied to accessibility and gene expression data alone, can recapitulate a “gold standard” network (Ciofani et al., 2012; Yosef et al., 2013) that was built through the more laborious approach of TF chromatin immunoprecipitation and TF KO RNA-seq, in related Th17 cells (Miraldi et al., 2018).

In this study, we applied our newest *Inferelator* method to construct TRNs for five intestinal ILC subsets (CCR6^+^ ILC3, CCR6^-^ ILC3, ILC1, NK cells, and ILC2) by leveraging recently published ILC genomics studies and our own intestinal ILC gene expression and chromatin accessibility datasets. This unbiased approach recovers the few known TF regulators of subset-specific genes and predicts new roles for nearly a hundred additional TFs in ILC regulation. Clustering of TFs, based on shared regulatory interactions, reveals core modules: groups of TFs that cooperatively regulate distinct functional pathways. Strikingly, the combined intestinal ILC network not only identifies activators of genes within each subset but also repressors of genes corresponding to alternative ILC lineages. We experimentally validate TFs predicted to repress alternative-lineage genes. These proof-of-concept validation experiments highlight how TRNs can be used to elucidate control points of cell identity and plasticity in mature intestinal ILCs.

## Results

### Innate lymphoid cell chromatin accessibility landscapes suggest activating and repressive regulatory elements

For construction of the transcriptional regulatory networks (TRNs) that underlie mature ILC maintenance and function, we globally identified putative regulatory elements in small (SI) and large intestine (LI) ILCs by the genome-wide assay for transposase-accessible chromatin using sequencing (ATAC-seq) (Buenrostro et al., 2013). In parallel, we also assessed gene expression by RNA-seq (Figure 1). Five ILC subsets were sort-purified from C57BL/6 mice using surface markers that discriminated lineage-restricted TF expression **(Figure S1)**. Canonical NK cells were negative for IL7rα and other lineage markers but expressed NK1.1 while T helper-like ILCs were positive for IL7rα and expressed either Klrg1 and Gata3 (ILC2) or Klrb1b (ILC1/ILC3 continuum). Expression of NK1.1 and CCR6 among Klrb1b+ cells further discriminated T-bet positive ILC1 from two ILC3 sub populations that either expressed RORγt alone (CCR6^+^, which includes LTi cells) or co-expressed RORgt and T-bet (CCR6^-^).

**Figure 1.**
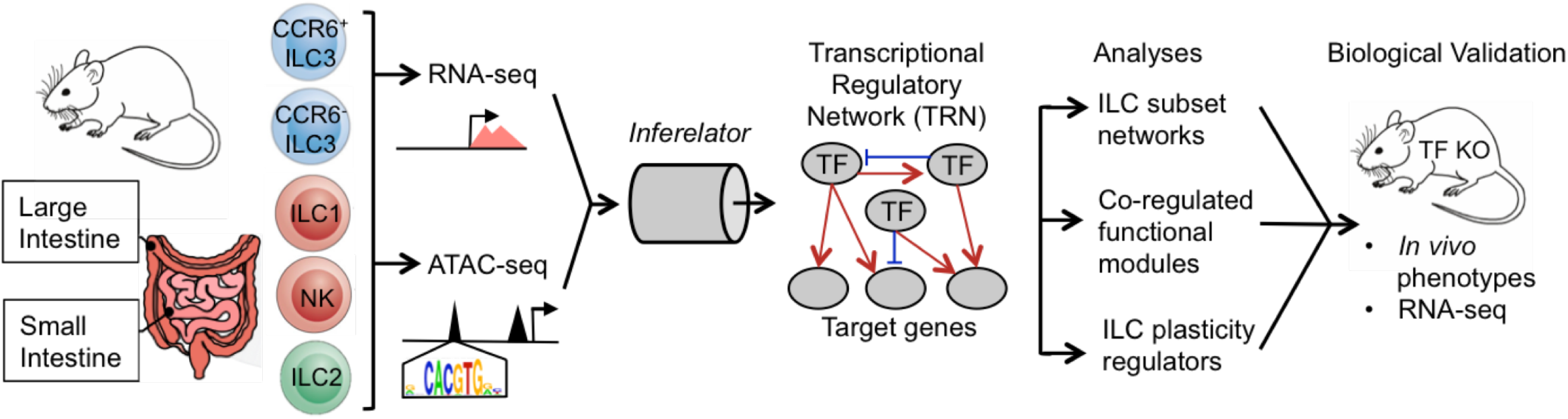
Study design for generation and biological validation of ILC TRN. ILCs were isolated from the small and large intestine of mice for measurement of gene expression (RNA-seq) and chromatin accessibility (ATAC-seq) to build a transcriptional regulatory network (TRN) explaining subset-specific gene expression of intestinal ILCs.

Quantitative analyses of the RNA-seq and ATAC-seq data led to the identification of differentially expressed genes and regulatory elements for each ILC subset. Correlation-based clustering of samples yielded expected groupings of ILCs based on the three broad ‘types’ with smaller differences between subsets, i.e., ILC1 and NK cells differed more from ILC2 and ILC3 than from each other (**Figure S2**). Examination of differentially expressed genes revealed the expected lineage-specific TF and cytokine/receptor genes (**Figure S3**). These include *Rorc, Il23r, Il22, Il1r1, Il1r2* in ILC3; *Gata3, Hes1, Pparg, Areg, Il1rl1* (ST2), *Bmp2, Bmp7* in ILC2; *Tbx21* in ILC1; and *Eomes* and *Gzma* in NK. In addition to these canonical ILC-subset markers, we identified hundreds of TFs and immunological signatures that also distinguished the subsets (**Figure S3**).

To explore major trends in the regulatory elements identified by ATAC-seq, we performed principal component analysis (PCA) on all (~127K) accessible peaks in the 5 ILC subsets (Figure 2A, left panel). The largest accessibility pattern distinguished ILC1/NKs from ILC3s (39% of the variance, PC 1), while the second largest separated ILC2s from ILC1s and ILC3s (27% of the variance, PC 2). Principal components 3 and 4 (not shown) captured smaller trends distinguishing ILC1 from NK and CCR6^+^ ILC3 from CCR6^-^ ILC3. Approximately 10^4^ peaks were unique to each broad class of ILC (ILC1/NK, ILC2, ILC3) (light colored circles, Loadings Plot, Figure 2A; light colored bars Figure 2B).

We next integrated differential gene expression into our analysis to determine if the putative regulatory elements correlated with expression of nearby genes. Peaks were assigned to genes based on their proximity (+/- 10kb) to genes uniquely expressed in one of the major ILC subsets (dark green, blue and red dots on the loadings plot Figure 2A, and dark colored bars Figure 2B). Indeed, the majority of proximal subset-specific peaks were adjacent to genes that were uniquely expressed in the same subset, consistent with putative promoter or enhancer activities (Figure 2B). One example is an ILC3-specific peak lying upstream of the *Il17re* locus, a gene that was uniquely expressed in ILC3s (Figure 2C, upper panel). Given the correlation between the gene expression and accessibility pattern, we posited that the accessible region might promote *Il17re* expression. A motif search in the ILC3-associated peak yielded a Retinoic Acid-Related Orphan Receptor Response Element (RORE), to which the ILC3-expressed TFs RORgt or RORa might bind and potentially promote *Il17re* expression.

Strikingly, a smaller number of ATAC-seq peaks proximal to genes uniquely expressed in ILC3 (dark blue dots and bars) were instead uniquely accessible in other subsets (ILC2s and ILC1/NKs, light green and pink) (Figure 2A,B). For example, two peaks, which were uniquely accessible in ILC1/NKs, are near a gene (*Il23r*) that is uniquely expressed in ILC3s (Figure 2C, lower panel). This pattern might arise from mechanisms of active repression. Specifically, NK/ILC1 lineage TFs might bind at distinct sites to suppress expression of alternative-lineage genes, such as *Il23r*. Based on TF gene expression patterns (**Figure S3**) and motif analysis, three potential ILC1/NK TFs (IRF8, TBX21 and EOMES) are candidate negative regulators of *Il23r* in type 1 ILCs. Similar to ILC3s, a smaller number of subset-specific peaks negatively correlate with the expression of proximal alternative-lineage genes in type 1 and 2 ILCs (Figure 2B).

To determine which TFs might bind to predicted activating or repressive regulatory elements in each ILC subset, we performed TF motif analysis (**Methods**). Specifically, we tested whether motifs of TFs uniquely expressed in a particular lineage were enriched in accessible regions proximal to genes expressed in that lineage (putative activation) or other lineages (putative repression). This analysis recovered motifs of the key known lineage-specifying TFs RORgt, T-bet and GATA3, as well as new candidate regulators (Figure 2D). For example, in unique ILC1/NK accessible regions, motifs for T-bet, Eomes, and other ILC1-expressed TF motifs were enriched proximal not only to ILC1 genes (suggesting activation) but also to ILC2 and ILC3 genes (suggesting that these TFs may have repressive functions across multiple gene loci).

**Figure 2.**
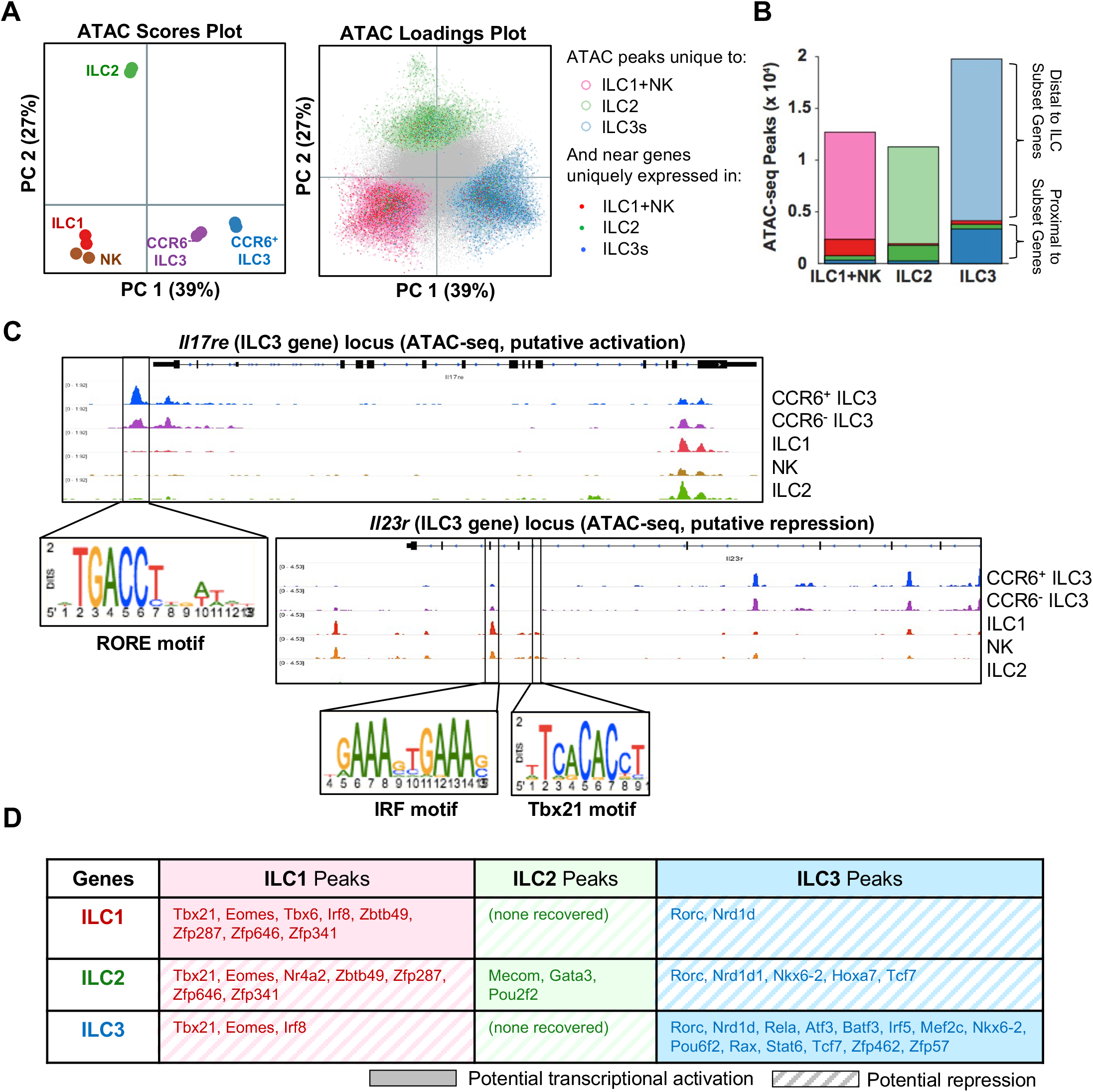
Characterization of genomic regulatory elements in ILCs. **(A)** 127k accessible regions (peaks) were identified in ILCs and accessibility levels quantified across samples. In the PCA scores plot, SI ILC samples are plotted according to accessibility patterns, while the PCA loadings plot highlights the peaks that contributed most to those patterns. In the loadings plot, peaks are annotated according to lineage-specific accessibility (pastel circles) and proximity to lineage-specific ILC genes (dark colored dots). (Lineage-specific peaks and genes are defined to have increased accessibility or expression in ILC1+NK, ILC2, or ILC3s at FDR=10%, log2(FC)>1 for each pairwise comparison.) **(B)** Subset-specific ATAC-seq peaks were categorized as proximal (+/-10kb gene body) or distal to subset-specific genes; color scheme as in (A). **(C)** Representative ATAC-seq data tracks are displayed for each ILC subset and TF motif occurrences in putative activating and repressive regions are highlighted in upper and lower panels, respectively. **(D)** The table displays subset-specific TFs whose motifs were enriched in subset-specific peaks proximal to subset-specific genes (FDR=10%, log2(FC)>1).

Thus, simple motif-enrichment analysis, integrating differential accessibility with gene expression patterns of peak-proximal genes, can be used to derive a noisy initial set of TF-gene regulatory interactions (Blatti et al., 2015) and can predict whether these interactions may be activating or repressive (Karwacz et al., 2017). However, such a regulatory network would be expected to contain many false positives and negatives. For example, sources of error would include: 1) A TF motif occurrence within a peak indicates the potential for, rather than evidence of, TF binding; 2) TF motifs are often shared by members of the same TF subfamily, leading to ambiguity; 3) long-range regulatory interactions are difficult to assign to genes in the absence of 3D-chromatin conformation data; and 4) not all TF motifs have been determined (leading to the exclusion of many potential regulators). Thus, we refer to this initial, noisy set of TF-gene interactions as a “prior” network (∆ATAC), serving as input to a more sophisticated method for TRN inference (Miraldi et al., 2018).

### ILC subsets have shared and unique core regulatory networks

We constructed an ILC TRN using our most recent *Inferelator* method (Miraldi et al., 2018) (Figure 1). Like the original (Bonneau et al., 2006), this method models gene expression as a function of TF activities, leveraging (partial) correlations between gene expression and TF activity profiles across many sample conditions to learn TF-target gene interactions (**Methods**). As increasing the number of gene expression samples boosts network inference, we incorporated publicly available ILC genomics datasets (Gury-BenAri et al., 2016; Shih et al., 2016) into our analysis (**Figure S4A**). As a result, our gene expression matrix tripled in size (to 62 ILC samples total: 40 from SI, 8 from LI, and the remainder from bone marrow, lung, liver and spleen). Importantly, recent versions of the *Inferelator* (Arrieta-Ortiz et al., 2015; Greenfield et al., 2013; Miraldi et al., 2018) enable incorporation of prior information (e.g., ∆ATAC prior) and have markedly improved inference in another mammalian context, Th17 cell differentiation (Miraldi et al., 2018).

The ILC TRN contains models for 6,418 genes, resulting in 63,832 TF-gene interactions mediated by 445 TFs across all subsets (detailed in **Methods**, **Figure S4**). TFs were ranked according to number of target genes in the network; fourteen of the top 100 TFs (e.g., ARNT2, IKZF2, FOXJ1) were not in the DATAC prior but had strong support from gene expression modeling alone, confirming that our ILC TRN was not limited to TFs with known motifs (**Figure S5**). Positive regulatory interactions outnumbered repressive interactions by 1.6:1 in the ILC TRN. This positive-edge bias was consistent with the distribution of lineage-specific ATAC-seq peaks, in which putative promoter elements were proximal to genes specific to that lineage more frequently than genes unique to another lineage (Figure 2B); this trend was mirrored in subset-specific TF-gene interactions (**Figure S4B,C**). Thus, the ILC TRN predicted both activating and repressive mechanisms driving subset-specific gene expression patterns.

To identify the key transcriptional regulators associated with each cell type, we derived “core” TRNs for each of the five ILC subsets. We identified TFs that strongly promoted the gene expression patterns of individual ILC subsets based on enrichment of (1) positive targets in the genes up-regulated in that subset or (2) repressed targets among down-regulated genes (Miraldi et al., 2018). This enabled recovery of both activating and repressive modalities for promotion of subset-specific gene expression patterns. TFs regulating subset gene signatures were clustered to identify classes of TFs unique to or shared between ILCs (Figure 3A). This analysis correctly recovered well-known subset-specific regulators. For example, RORgt is recovered as a shared regulator of both ILC3 subsets (Sawa et al., 2010). GATA3 is recovered in the ILC2 network (Hoyler et al., 2012). T-bet is a shared regulator in the NK and ILC1 networks (Powell et al., 2012; Sciumé et al., 2012; Townsend et al., 2004), while Eomes is unique to NK (Pikovskaya et al., 2016). Beyond these lineage-defining TFs, the ILC networks accurately predicted several other TFs with known functions in various ILC lineages. These include TCF-1 (*Tcf7*) in CCR6^+^ ILC3 (Mielke et al., 2013); LEF1 (Held et al., 2003) and BLIMP-1 (*Prdm1*) (Kallies et al., 2011; Smith et al., 2010) in NK; KLF2 in NK and ILC1 (Rabacal et al., 2016); and PPARg in ILC2 (highlighted in bold, Figure 3A). Importantly, the majority of the 125 TFs are novel predictions, highlighting the utility of our network approach to markedly expand TF predictions beyond what is possible by standard motif enrichment analysis.

**Figure 3.**
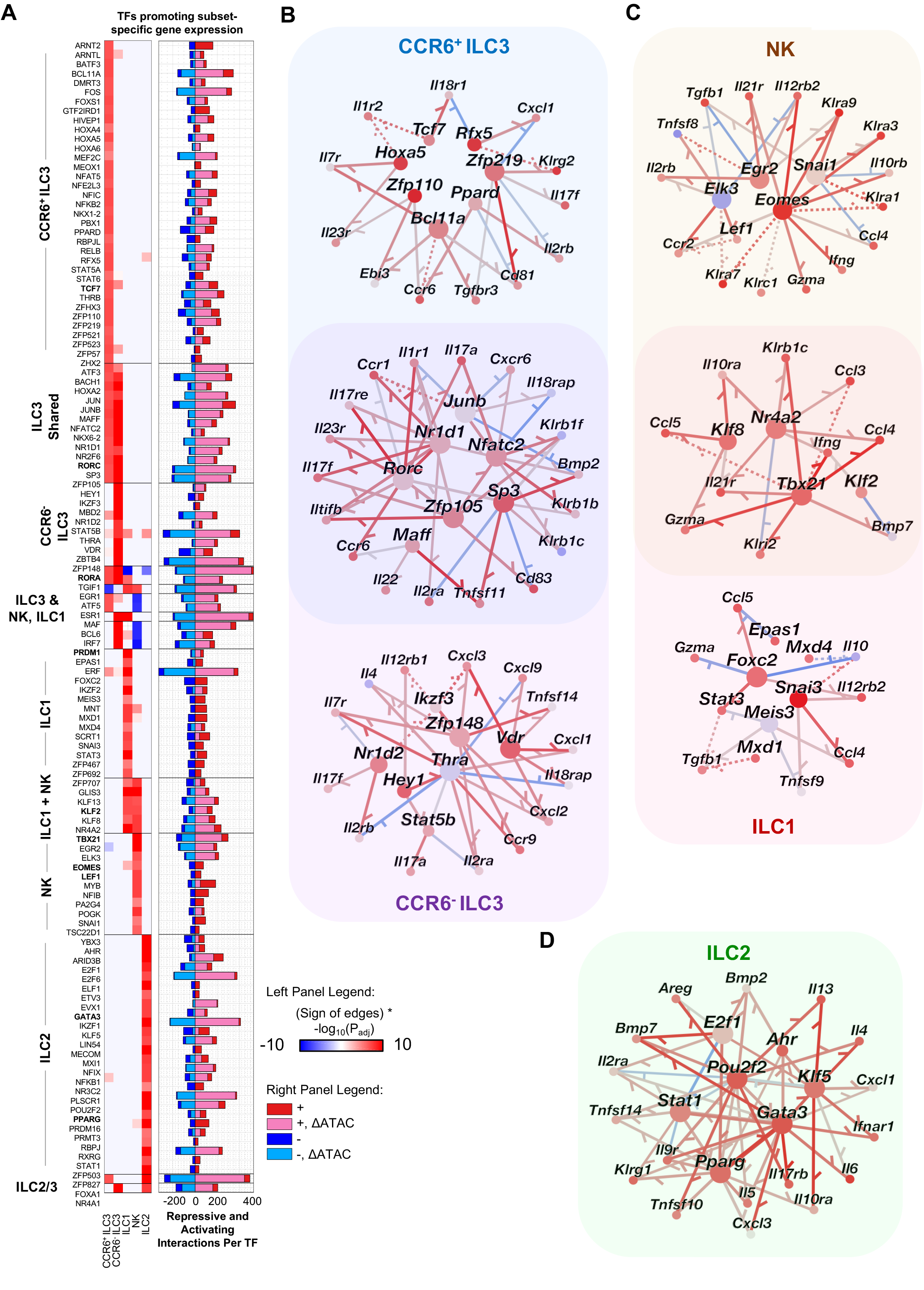
Core TRNs for the ILC subsets. **(A)** Core TFs are clustered according to enriched positive regulation of up-regulated genes (red) or negative regulation of down-regulated genes (blue) in a given ILC subset, as both types of regulation would contribute to subset-specific gene expression. The significance of enrichment is displayed in the left panel. The right panel displays the number of positive (red) and negative (blue) regulatory interactions per TF; interactions supported by the DATAC prior are shaded pink (+) and blue (-). TFs previously associated with a given ILC subset are bolded. Core TRNs are displayed for **(B)** ILC3, **(C)** ILC1, and **(D)** ILC2 subsets, representing subset-specific as well as shared regulatory modalities, for select TFs in (A). Target genes are limited to differentially expressed cytokines, chemokines, and receptors. Node color represents z-scored gene expression in the given cell type (red for up-regulation and blue for down-regulation); TF node size is determined by its degree (number of target genes). Edge color denotes positive (red) and negative (blue) regulation. Solid edges are supported by the ∆ATAC prior and gene expression modeling, while dotted edges are supported by gene expression modeling alone.

To examine what genes were regulated by core TFs, we visualized the subset-specific TRNs, limiting target genes to cytokines and receptors (Figure 3B). Several of the cytokine interactions have been previously verified. For example, RORγt in ILC3 was predicted to regulate its canonical target genes, including *Il23r*, *Il17a*, and *Il1r1*. Thus, the recovery of known TFs and their regulatory interactions provided confidence in the overall quality of the networks and their value for pursuing novel interactions. These searchable core TRNs are available to explore via an interactive web-based interface: https://github.com/flatironinstitute/ILCnetworks (see **Supplemental Information** for instructions).

### Transcription factor co-regulation of distinct functional pathways

The analyses described above implicated individual TFs in transcriptional control of genes specific to each ILC subset. However, many TFs function in a combinatorial manner to co-regulate distinct gene programs that enable specific cellular functions (Pope et Medzhitov, 2018). To determine how the TFs in the ILC TRN connect to ILC functions we clustered TFs into “co-regulating modules” based on overlap in their target genes (Miraldi et al., 2018). This analysis identified several groups of TFs with significant overlap in their positive target genes and helped organize the network into easier-to-interpret modules (Figure 4A, **Methods**). To help identify functional roles for each module, we performed gene-set enrichment analysis on the module’s co-regulated target genes (**Figure S6**, representative gene ontology and cellular pathway enrichments are summarized in Figure 4A). A cluster composed of TGIF, NR2F6, IRF7, VDR, MAF, and BCL6 is predicted to positively regulate chemotaxis and cytokine signaling, suggesting that it may be relevant to ILC identity and trafficking. Each of these factors, with the exception of TGIF1, were predicted to be both activators and repressors of ILC3 genes. These TFs are most highly expressed in SI CCR6^-^ ILC3, and are hence most likely to coordinate cytokine and chemokine programs in these cells (Figure 4B). The targets of known circadian regulators NR1D1, RORA, and NR1D2 (Sato et al., 2004; Zhang et al., 2014) are most highly expressed in ILC3, and form another distinct cluster (Figure 4A). A cluster of genes predicted to regulate adipogenesis is most highly expressed in ILC2 (in both SI and LI), and two of the TFs (GATA3 and STAT1) are themselves in the adipogenesis gene set (Figure 4A,C). ILC3s have the ability to process and present antigen to regulate T cells in response to microbiota (Hepworth et al., 2013, 2015), yet the transcriptional regulation of this process is not fully understood. This analysis predicted that a set of TFs (ZFP385A, SPI1, POU2AF1, and GATA2), most highly expressed in CCR6^-^ ILC3 of the LI, regulate genes involved in “Antigen presentation”, “Asthma”, “Rheumatoid Arthritis”, and “Intestinal IgA production network”. Indeed, several HLA genes are regulated by TFs in this module, including *H2-DMa*, *H2-DMb2*, *H2-Ab1*, *H2-DMb1*, and *H2-Ob* (Figure 4D). Thus, our coregulator module analysis highlights sets of TFs with coordinated regulation of key pathways in ILCs, recovering some known TF-pathway associations and many more novel predictions for TF regulation of ILC function.

**Figure 4.**
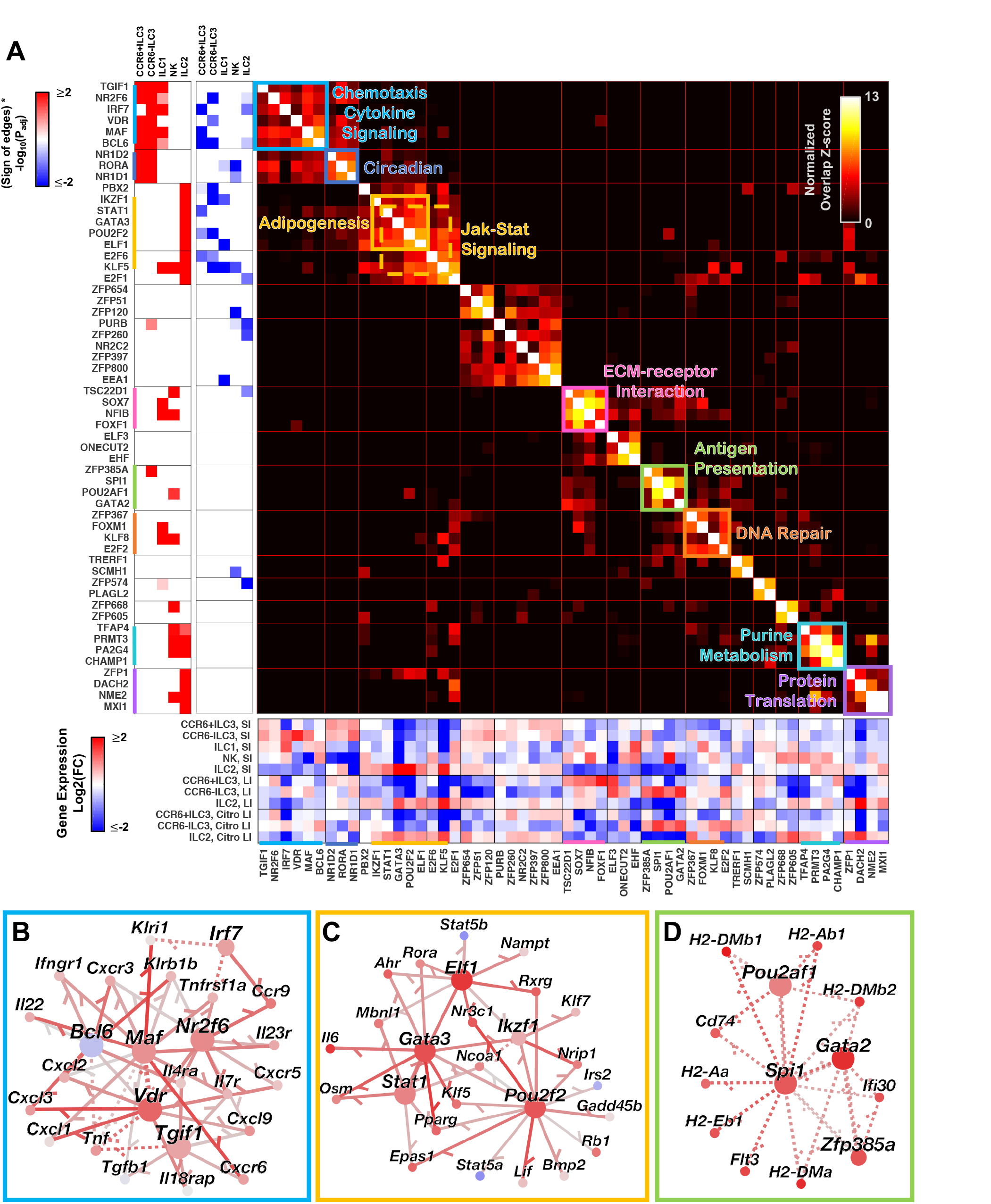
TF-TF modules co-regulate expression of gene pathways. **(A)** The normalized target gene overlap between the top 15 “TF-TF modules” (sets of TFs with significant overlap in positively regulated target genes) are displayed and annotated with enriched gene pathways in the main panel. TF-TF module positive and negative target gene enrichments in ILC subset gene sets are displayed in the left panels, while gene expression of TF constituents is displayed in the bottom panel. Positive TFTF modules are displayed for the **(B)** “chemotaxis, cytokine signaling”, **(C)** “adipogenesis”, and **(D)** “antigen presentation” modules, and nodes represent z-scored relative gene expression in SI CCR6-ILC3, SI ILC2, and LI CCR6^-^ ILC3, respectively. Target genes were included in the TRN visualization if positively regulated by at least two TFs in the module.

### The ILC TRN reveals TF repressors of alternate lineage genes

The phenomenon of ILC functional and phenotypic plasticity is poorly understood. We hypothesized that regulatory circuits directing the conversion of one ILC subset into another might function via de-repression of alternative ILC subset genes. We thus focused on TFs in the network that were predicted to act as suppressors of such genes. We identified TFs in the TRN whose negative target genes were enriched in signature genes of another ILC subset (Figure 5A). Several of the TFs (TBX21, EOMES, IRF8) identified by DATAC motif enrichment analysis (Figure 2D) were again recovered by this TRN analysis. Importantly, the TRN also uncovered numerous additional factors, several of which did not have recognizable TF motifs and were thus unrecoverable from ATAC-seq analysis alone. We further examined how these negative regulators were predicted to regulate one another. We thus generated a TF-TF lineage-regulating network for some of the most significant alternative-lineage gene repressors identified (Figure 5B). In the repressive TRN, ILC3-specific *Rorc* antagonizes ILC1/NK regulators *Tbx21* and *Bach2*, while NK/ILC1 regulators, including *Tbx21* and *Bcl6*, repress *Rorc*. *Tbx21* additionally antagonizes the hallmark ILC2 TF *Gata3*. Reciprocally, ILC2 TFs *Klf5*, *Pou2f2* and *Irf4* repress *Tbx21* and *Rorc*. These analyses implicate a number of ILC regulators in the control of ILC plasticity. We chose BCL6 and c-MAF as candidates for further validation due to their predicted reciprocal regulation of genes in the ILC1/ILC3 continuum as well as for their predicted roles in controlling a module of cytokine and cytokine receptor genes (Figure 4A,B).

### c-MAF regulates the balance between intestinal ILC1 and ILC3

The transcription factor c-MAF has critical functions in effector T cells, contributing to the expression of IL-10 in multiple T cell subsets, the differentiation of induced T_reg_ cells (Xu et al., 2018), and the repression of IL-22 transcription (Rutz et al., 2011). Its function in ILCs has not been explored. Our ILC TRN predicted that c-MAF could repress genes unique to the CCR6^+^ ILC3 lineage and promote expression of genes in the CCR6^-^ ILC3 and ILC1 lineages, where c-MAF is most highly expressed (Figures 3A, 4A, 5A). To address whether c-MAF regulates these cell types, we examined ILCs in the SI lamina propria of mice lacking c-MAF in all lymphocyte populations (*Il7ra^cre^*; *Maf^fl/fl^*) (Wende et al.,2012) (Figure 6A). Indeed, there was a reduction in the number of RORγt^+^ T-bet^-^ ILC3 accompanied by an increase in the frequencies and total numbers of the other ILC subsets, particularly ILC1, in *Maf^fl/fl^;Il7ra^cre^* animals compared to *Maf^fl/fl^* (or *Il7ra^cre^*) controls (Figure 6A,C). Notably, *Maf^fl/fl^;Il7ra^cre^* ILCs exhibited increased expression of T-bet, suggesting that c-MAF may regulate the balance of ILC1 and CCR6^+^ ILC3 via direct or indirect suppression of *Tbx21* (Figure 6B). Thus, as predicted by the ILC TRN, c-MAF regulates the balance between ILC1 and ILC3.

### BCL6 promotes abundance of intestinal type 1 ILCs

The ILC TRN predicted that BCL6 was a core regulator of NK and CCR6^-^ ILC3 lineages (Figure 3A). BCL6 is a transcription factor critical for controlling germinal center B cell and follicular helper T (Tfh) cell differentiation (Hatzi et al., 2013, 2015), with a known role for alterative-lineage repression of Th17, Th2 and Th1 programs in Tfh (Hatzi et al., 2015). However, a role for BCL6 in ILC function has not been previously characterized. Conditional deletion of BCL6 in NK, ILC1 and NKp46-expressing ILC3 was achieved by crossing *Bcl6^fl/fl^* (Hollister et al., 2013) to *Ncr1^Cre^* (Narni-Mancinelli et al., 2011) mice. Comparison of SI lamina propria ILC populations between *Bcl6^fl/fl^; Ncr1^Cre^* (*Bcl6* KO) and control *Bcl6^fl/fl^* mice (or *Ncr1^Cre^* mice) revealed a decrease in NK and ILC1 cell frequencies and numbers (Figure 6D, E, F). There was no change in the frequency or numbers of ILC3 and ILC2. *Bcl6* expression was highest in NK and then decreased progressively from ILC1 to CCR6^-^ ILC3 to CCR6^+^ ILC3 (**Figure S3A, Table S3**). Thus, we observed a decrease in the two ILC populations with highest *Bcl6* expression. Intriguingly, the ILC TRN additionally predicted that BCL6 both positively and negatively regulated ILC3 signature genes (Figures 3A, 5A). Thus we sought to investigate BCL6-dependent gene changes to test the accuracy of target gene prediction by the *Inferelator* and to shed light on how BCL6 regulates ILC homeostasis and function.

**Figure 5.**
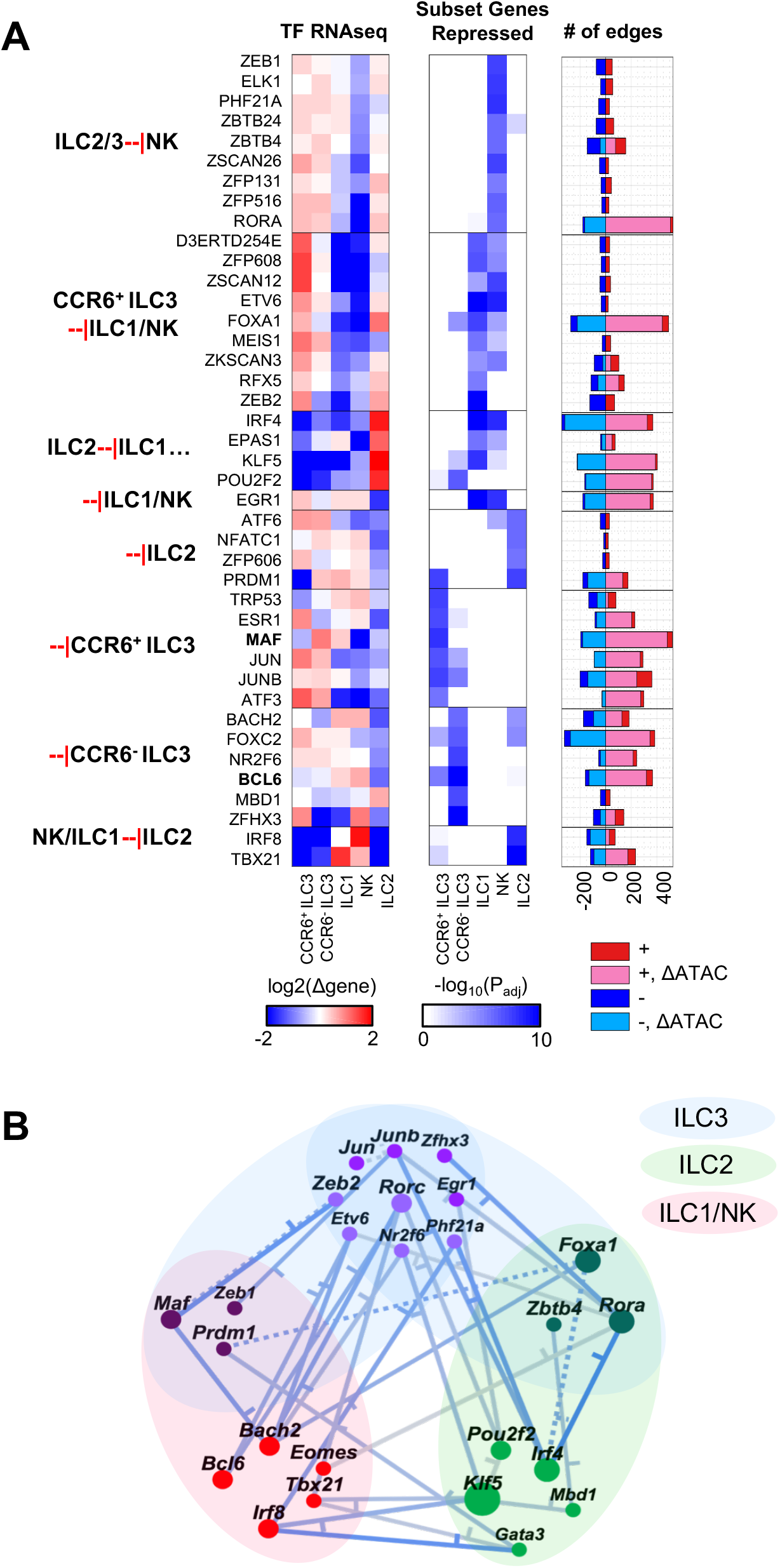
Predicted TF repressors of alternative ILC subset genes. **(A)** TFs predicted to be repressors of subset-specific gene expression patterns are clustered according to the significance of negative edge enrichment in genes upregulated per subset (FDR=1E-5; blue heatmap, middle panel). The left panel displays relative gene expression for the ILC subsets in the SI, while the right panel indicates number of target genes per TF, as in Figure 2A. **(B)** A TRN containing only TFs was constructed from a selection of lineage-repressors in addition to the “master regulators” RORC and GATA3. Nodes are colored according to ILC subset.

**Figure 6.**
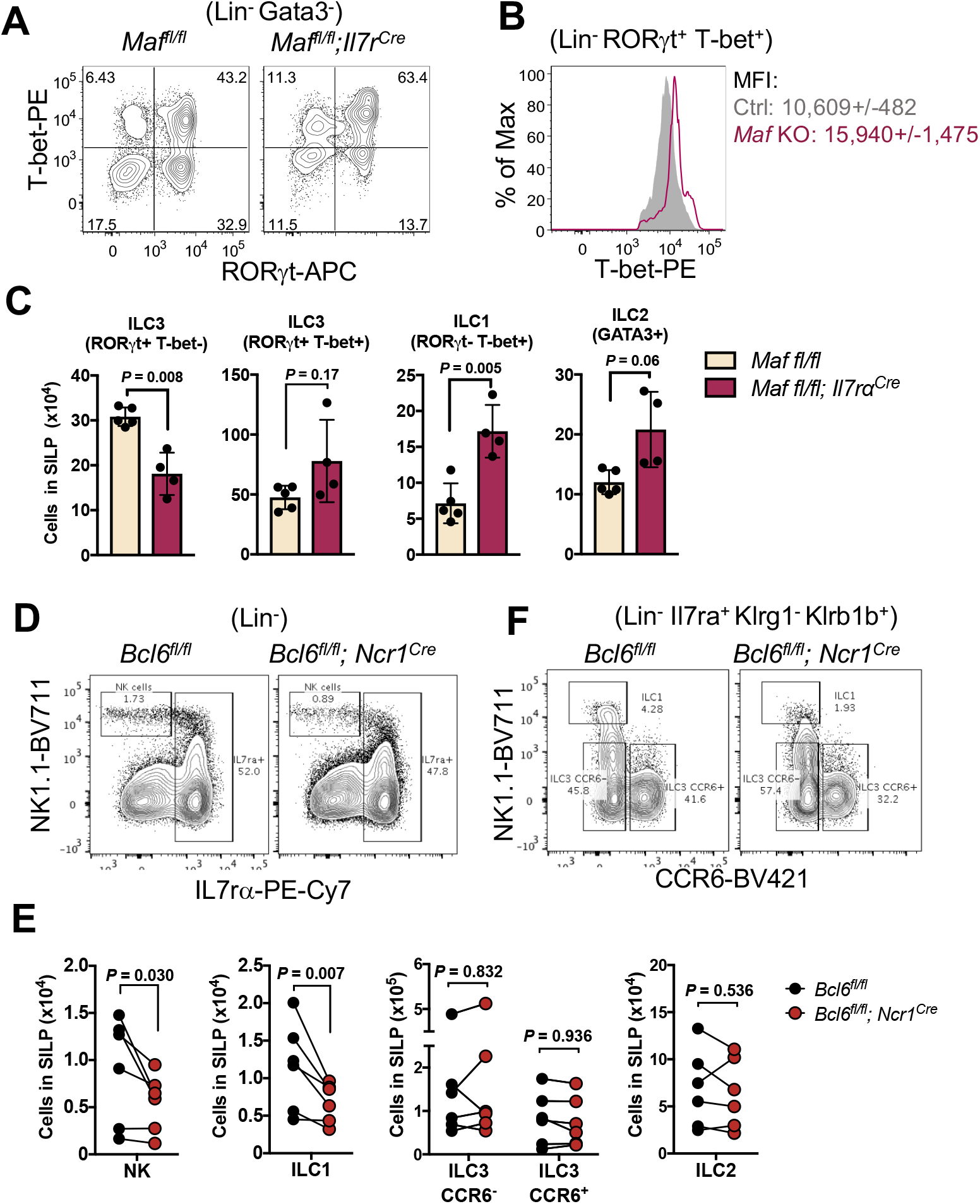
Functional roles for c-MAF and BCL6 in intestinal ILCs. **(A)** Indicated ILC populations from the SILP of *Maf^fl/fl^;Il7r^Cre^* or control *Maf^fl/fl^* mice. Lineage negative is (CD3^-^TCRb^-^TCRgd^-^CD11b-CD11c^-^CD14^-^CD19^-^). **(B)** Histogram depicting T-bet expression in RORgt^+^T-bet^+^ ILC3 gate. MFI is mean fluorescence intensity +/- standard deviation. **(C)** Quantification of absolute numbers of indicated ILC populations from one of three representative experiments. Statistics calculated by Welch’s *t* test. **(D, E)** Indicated ILC populations from the SILP of *Bcl6^fl/fl^; Ncr1^Cre^* or control *Bcl6^fl/fl^* mice. Lineage negative is as in (A). **(F)** Quantification of absolute numbers of indicated SILP ILC populations. Each line corresponds to an independent experiment (n=6) containing between 2-4 mice of each genotype. Each dot represents mean value for each experiment. Statistics calculated by ratio paired t test.

### Validation of BCL6 as a modulator of NK cell and ILC3 gene programs

To gain further insight into the regulatory footprint of BCL6 in the ILC lineages, we compared transcriptomes of NK, ILC1, and CCR6^-^ ILC3 from *Bcl6* KO and control mice. *Bcl6* deletion led to gene expression changes in each subset with greatest effects in NK cells, resulting in nearly 1000 differentially expressed genes (FDR = 25%), composed of a near-equal number of up-regulated and down-regulated genes (Figure 7A). Although less highly expressed in these cells, deletion of *Bcl6* also significantly perturbed expression patterns in CCR6^-^ ILC3, leading to 800 differentially expressed genes. The majority of the *Bcl6*-dependent genes in CCR6^-^ ILC3 were up-regulated, suggesting a largely repressive role for BCL6 in ILC3 (Figure 7A). Although SI ILC1 have the second-highest level of *Bcl6* expression (**Figure S3A**), perturbation of *Bcl6* in ILC1 led to the smallest number of gene expression changes (fewer than 300).

**Figure 7.**
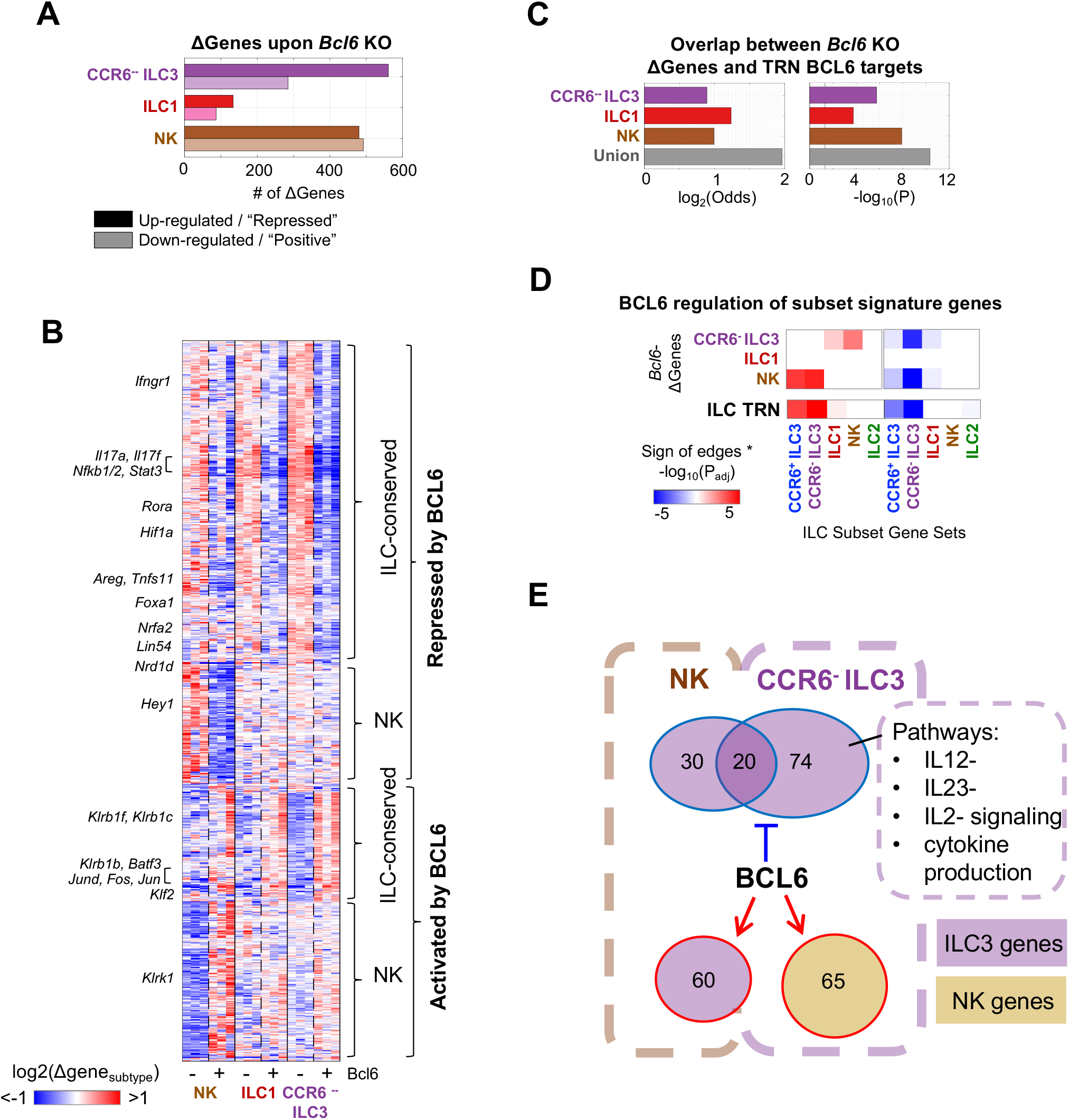
Validation of TRN predictions for BCL6. **(A)** The number of up-regulated and down-regulated genes comparing *Bcl6* KO (*Bcl6^fl/fl^ -Ncr1*^Cre*+/-*^) to control (*Bcl6^fl/fl^*) per ILC subset in the SI (FDR=25%). **(B)** Hierarchical clustering of 902 differentially expressed genes (*Bcl6* KO versus control (FDR=10%), in at least one of the three ILC subsets). Genes are normalized independently for each subset to highlight *Bcl6*-dependent trends. **(C)** Significance of overlap between TRN-predicted BCL6 targets and the *Bcl6*-KO-dependent genes (TRN BCL6 targets were >2-fold more likely to be differentially expressed; Fisher Exact Test P<1E-3, all subsets, and P<1E-10 for union of differential genes across subsets). **(D)** Positively and negatively-regulated BCL6 targets were (1) inferred from the *Bcl6* KO differential expression analysis in each cell type as well as the union across cell types (FDR=10%) and (2) tested for enrichment in genes upregulated per subset; TRN predictions for BCL6 (from Figure 4A) are provided for reference. **(E)** BCL6 was a significant regulator of lineage genes in NK cells and CCR6^-^ ILC3. GSEA of *Bcl6-*repressed targets in *Bcl6* KO CCR6^-^ ILC3 revealed enrichment of several gene pathways.

We clustered the *Bcl6* KO data to visualize overlap in differentially expressed genes across the subsets (Figure 7B) and observed several key trends. Across the four subsets, BCL6 has a greater number of putative repressed targets than positive targets consistent with its known roles as a transcriptional repressor (Chang et al., 1996; Crotty et al., 2010; Hatzi et al., 2015) (Figure 7B). Among both repressed and positive targets, genes cluster into those that are conserved across NK, ILC1, and ILC3 and those that are unique to NK cells. Most of the putative repressed targets (70%) are conserved among ILCs and notably include canonical ILC3 genes such as *Rora, Stat3, Il17a, Il17f*, while 67% of positive targets are NK-cell-specific (Figure 7B). It should be noted that, although fewer genes were significantly differentially expressed in ILC1 (Figure 7A), Bcl6-dependent gene patterns in ILC1 were qualitatively similar to CCR6^-^ ILC3, suggesting conservation of regulatory mechanisms (Figure 7B). Genes that appear to be positively regulated by BCL6 include several with known roles in NK cell differentiation and functions (*Klf2, Fasl, Clec2d, CD48, CD160, Ptpn6, Klrk1, Klrb1b, Klrb1c, Klrb1c, Klrb1f*). These results suggest that in ILC1 and ILC3, BCL6 is mainly a repressor, while, in NK cells, BCL6 is both a repressor and an activator (Figure 7B, **S7C,D**).

To assess the accuracy of network predictions, we tested how well ILC TRN-predicted BCL6 targets matched *Bcl6* KO gene expression data. Indeed, for each ILC subset, TRN-predicted targets were two-fold more likely to be differentially expressed in *Bcl6*-deleted cells (P<1E-4, Fisher Exact Test, Figure 7C), and the enrichment grew to four-fold upon combining differential genes across the ILC subsets (“Union”) (P<1E-10). These results were robust over a range of FDR cutoffs and TRN model sizes (**Figure S7A**). Importantly, TRNs built from gene expression data alone (without ATAC-seq data) were poor at predicting *Bcl6*-dependent genes (**Figure S7B**), highlighting the importance of integrating ATAC-seq data into TRN inference for prediction of BCL6 targets.

We next asked what genes BCL6 regulated and in which direction (Figure 7D). Strikingly, both up-regulated and down-regulated gene sets in *Bcl6* KO NK cells were enriched for ILC3-signature genes, supporting the ILC TRN prediction that BCL6 is both a positive and negative regulator of ILC3 genes in NK cells (Figure 7D). In CCR6^-^ ILC3, BCL6 also tuned down a subset of CCR6^-^ ILC3 signature genes, but too a lesser degree than in NK, perhaps due to its lower level of expression there. The CCR6^-^ ILC3 *Bcl6* KO data additionally suggested that BCL6 is a positive regulator of ILC1/NK-signature genes; some promotion of ILC1/NK-signature genes was observed in NK cells and ILC1s as well (**Figure S8A**). This result was not predicted by analysis of the ILC TRN and highlights the value of ILC subset-specific TF perturbation data to tease out additional targets of BCL6. There was significant overlap between BCL6-repressed CCR6^-^ ILC3 genes across NK cells and CCR6^-^ ILC3, while the BCL6-activated CCR6^-^ILC3 genes in NK do not overlap with the BCL6-repressed genes in any of the subsets (Figure 7E). Among TFs suppressed by BCL6, RORA (highest expression in ILC2 and ILC3) was also predicted to repress NK/ILC1 signature genes (Figure 5), highlighting a potential hypothesis to explain increased ILC1/NK gene expression in *Bcl6* KO ILCs.

BCL6 (and c-MAF) belong to a cluster of TFs predicted to co-regulate cytokine signaling (Figure 4B). Gene-set enrichment analyses on BCL6 targets from *Bcl6* KO confirmed that, in ILC3, BCL6 negatively regulated IL12-, IL23- and IL-2-signaling pathways, cytokine production, and endogenous TLR signaling (FDR<1%) (Figure 7E, **S8B**. Taken together, these results highlight the utility of the *Inferelator* in predicting gene regulation by validating activated and repressed target genes of BCL6 in ILCs. These analyses confirm a role for BCL6 as a modulator of the ILC3 program and activator of NK genes, with distinct roles in ILC3 and NK cells. These data suggest potential mechanisms whereby BCL6 maintains SI NK/ILC1 cells through repression of alternative lineage (ILC3) genes and promotion of select genes required for maintenance of these lineages.

## Discussion

Innate lymphoid cells are important mediators of mucosal immunity, especially in early immune responses. However, they also contribute to diverse pathologies, from autoimmune disease to metabolic syndrome. Previous ILC genomics studies identified genes, gene pathways and putative regulomes of the ILC lineages (Gury-BenAri et al., 2016; Koues et al., 2016; Robinette et al., 2015; Shih et al., 2016). In this study, we integrate our own substantial ILC genomics dataset with previous work (Gury-BenAri et al., 2016; Koues et al., 2016) to reconstruct the first genome-scale transcriptional regulatory network (TRN) for gut-resident ILCs. This effort also represents the first application of our TRN inference method (Miraldi et al., 2018) *in vivo*, providing an important proof of concept. The ILC TRN *de novo* predicted known critical regulators of ILC lineage as well as many new candidate TFs, two of which, BCL6 and c-MAF, we have confirmed to be functionally relevant in the contexts predicted. Moreover, the networks identify specific gene targets of each TF, further facilitating biological hypothesis development and testing. This TRN and co-regulated gene modules are intended to serve as a resource for the ILC field. Networks and gene expression data are available via a user-friendly web interface that enables searchable interactive visualization of both networks and gene expression.

By integrating gut ILC subsets into a single network, we were able to explore regulatory relationships between as well as within ILC subsets. One of the striking findings from analysis of the combined ILC TRN was an extensive chromatin landscape of alternative lineage gene repression. Many of the repressive interactions were supported by ATAC-seq peaks with high accessibility and binding motifs for TFs expressed within a particular subset but adjacent to alternative lineage genes that were not expressed. One regulatory element in the *Ifng* gene with this pattern of accessibility in ILC2 was highlighted previously (Koues et al., 2016). Our work demonstrates that this regulatory pattern is broadly present within all ILC subsets and may represent a general mechanism underpinning ILC lineage regulation. Such elements may function as silencers and, together with poised and active enhancers, respond to environmental cues to modulate ILC identity.

For each of the five ILC subsets in our study, we defined “core” TFs and TRNs, recovering known TF lineage regulators and implicating nearly 100 additional TFs in the regulation of lineage-specific gene expression patterns. Additionally, we detected over 50 TFs whose repressed targets in the TRN were enriched in lineage-specific genes. These TFs represent candidate lineage control points. Not only could they be key to maintenance of subset-specific gene expression patterns, but their perturbation could potentially be used to alter ILC lineage identity and functionality. Several autoimmune and inflammatory diseases are associated with altered abundances and activity of ILC subsets (Eberl et al., 2015). For example, maternal antibiotic exposure is associated with a decrease in lung ILC3s and an increased susceptibility to infection (Gray et al., 2017). Thus, our ILC TRN could be used to design disease interventions that limit the function or abundances of specific ILC subsets through targeting of lineage-promoting or lineage-repressing TFs. Construction of ILC TRNs using human ILCs derived from disease contexts could provide additional clues as to what TFs are active in these states. We also emphasize that our approach is entirely general and readily applicable to other plastic cell populations in physiological settings, including T helper cells, macrophages, and others.

We functionally tested predictions from our “core” TRN and alternate-lineage repressor analyses. We predicted that MAF was a “core” positive regulator of lineage-specific gene programs in ILC1 and CCR6^-^ ILC3. We also predicted that MAF suppressed the CCR6^+^ ILC3 program. Thus, we anticipated that deletion of MAF would shift the ILC3-ILC1 continuum. Indeed, these populations were altered in *Maf^fl/fl^ Il7r^Cre^* KO mice relative to control. We also tested the predicted role of BCL6 as a “core” regulator of NK and CCR6^-^ ILC3 lineages using conditional *Bcl6* KO mice. *Bcl6* deletion led to a decrease in SILP NK and ILC1, while there was no change in CCR6^-^ ILC3 frequencies.

By profiling gene expression in the three *Bcl6*-expressing ILC populations of the SI, we confirmed that BCL6 was both a positive and negative regulator of ILC3 genes. The subset-specific *Bcl6* KO data was critical to confirming TRN BCL6 target predictions and pinpointing the specific ILC subsets in which BCL6 positively and/or negatively regulated ILC3 and other target genes. BCL6 was found to significantly antagonize the ILC3 program in NK and CCR6^-^ILC3, and less so in ILC1. The *Bcl6* KO gene expression experiments and their enrichment analyses provide a potential explanation for the reduction of SI ILC1 in *Bcl6* KO animals. BCL6 might contribute to ILC3-to-ILC1 plasticity through repression of ILC3-promoting signaling pathways (e.g., IL23 signaling). BCL6 also positively contributes to NK and ILC1 gene expression (e.g. *Klf2*), and this represents a second mechanism by which BCL6 might contribute to maintaining SI type 1 ILCs via promotion of genes required for survival or homing. We speculate that these two mechanisms may in fact be connected as BCL6 may repress ILC3 TFs (e.g. RORA) in type 1 ILCs that in turn repress type 1 signature genes affecting ILC1 and NK cell accumulation within the intestines. Further experimental evidence is required to explore which BCL6 target genes enumerated by the ILC TRN are critical for the observed phenotypes.

Notably, BCL6-repression of the ILC3 program mirrors BCL6 repression of the Th17 program in Tfh cells. For example, in both ILCs and Tfh, *Rora*, *Stat3*, *Il17a*, *Il17f* are BCL6 targets while *Rorc* is not (Hatzi et al., 2015). Thus, like other TFs (Walker et al., 2013), BCL6 shares some regulatory roles between ILC and T Helper subsets. Application of our methods to construct TRNs of intestinal T helper cell TRNs would provide an opportunity for a comparative analysis of shared and unique regulatory mechanisms in T helper cells and ILCs.

In NK cells, the purpose of BCL6 repression of ILC3 genes is less clear, as plasticity between NK and ILC3 (or NK and ILC1) has not been reported. We note that our study is the first to contribute both NK and ILC1 RNA-seq and ATAC-seq data from the intestine, and to show that these two subsets have stronger correlation in accessibility patterns than previous studies comparing intestinal helper ILC subsets to spleen and bone marrow NK cells (Koues et al., 2016). As another potential explanation, NK cells might need to express ILC3 signature genes under some contexts.

A number of ILC TRN predictions are consistent with current knowledge of ILC lineage regulation, and a limited number of novel predictions have been experimentally tested in this study. However, the vast majority of predictions remain to be tested. We anticipate that the ILC TRNs will be useful to unraveling the molecular mechanisms driving complex phenotypes (e.g., due to TF KO or other experimental perturbations) in ILCs. We hope that these maps of ILC lineage-specific transcriptional regulation increase the pace of discovery in ILC biology, eventually leading to the design of precise genetic and chemical perturbation of subset-specific ILC behaviors in the context of human health and disease.

## Acknowledgements

We thank the Flatiron Institute Scientific Computing Core (I. Fisk) for enabling the computational aspects of this work and the New York University Langone Medical Center Genomics Core (A. Heguy and P. Zappile) for help with sequencing. This work was supported by the Cincinnati Children’s Research Foundation (Trustee Award to E.R.M., Research Innovation Award to S.N.W.; Research In Residency Award to N.S.C.; and Arnold P. Strauss Fellowship to D.O.), the Simons Foundation (E.R.M., A.W., N.D., N.C., R.B.), U.S. National Institute of Health (5T32AI100853 to M.P.; R01-DK103358-01 to R.B. and D.R.L.; DA038017 to S.N.W.; and R01-GM112192-01 to R.B., T32 CA009161 (Levy) to J.A.H.), Damon Runyon Cancer Research Foundation (Dale and Betty Frey Fellowship to J.A.H.), the Laura and Isaac Perlmutter Cancer Center (P30CA016087 to A.H.), and the Colton Center for Autoimmunity (D.R.L). Cell sorting and flow cytometric data acquired at Cincinnati Children’s relied on equipment maintained by the Research Flow Cytometry Core, which is supported in part by NIH grants AR47363, DK78392 and DK90971.

## Author Contributions

ERM, MP, JAH, DRL, and RB designed and contributed to analysis throughout the study. MP and JAH generated ILC genomics data. MP contributed c-MAF data. SNW conceived of BCL6 experiments and generated the mouse model. ERM, MP, JAH, DEO, SNW, DRL designed BCL6 experiments. DEO, NSC, HS performed BCL6 experiments. ERM built the ILC TRNs and computational analyses. RY compiled public ILC genomics data. NC mapped RNA-seq and ATAC-seq data. AW designed interactive network visualization software with input from ERM, MP and RB. ERM and MP wrote the manuscript with input from JAH, DEO, SNW, DRL, and RB.

## Declarations

The authors declare no competing interests.

## Methods

### Mice

*Wild type and Maf KO experiments. Maf^fl/fl^* and *Il7ra^Cre^* mice were kindly provided by C.Birchmeier and H. R. Rodewald (Schlenner et al., 2010; Wende et al., 2012). *Maf* conditional KO mice were generated by crossing *Maf^fl/fl^* to *Il7ra^Cre^* animals. *Maf* KO mice were bred and maintained in the animal facility of the Skirball Institute (New York University School of Medicine) in SPF conditions. All animal procedures were performed in accordance with protocols approved by the Institutional Animal Care and Usage Committee of New York University School of Medicine.

*Bcl6 KO experiments*. Mice were housed under pathogen-free conditions, and experiments were performed using ethical guidelines approved by the Institutional Animal Use and Care Committees of Cincinnati Children’s Hospital Medical Center. Ncr1^iCre^ mice were previously described (Narni-Mancinelli et al., 2011) and were kindly provided by Eric Viviér (University of Marseille). Bcl6^fl^/^fl^ mice were purchased from Jackson laboratories (Bar Harbor, ME). Both strains of mice were on C57BL/6 background. Ncr1^iCre^/^iCre^ mice and Bcl6^fl^/^fl^ mice were bred to obtain Bcl6^fl^/^fl^ Ncr1^wt^/^iCre^ (Bcl6 KO) or Bcl6^fl^/^fl^ Ncr1^wt^/^wt^ (WT). After breeding, all mice were kept with their littermates (males and females were separated). Bcl6 deletion was verified by examining NCR1 expression and by PCR. Male mice at 16–20 weeks of age were utilized in experiments. In each experiment, an equal number of mice was used.

### Isolation of SILP and LILP lymphocytes and TF analysis

Intestinal tissues were sequentially treated with PBS containing 1 mM DTT at room temperature for 10 min, and 5 mM EDTA at 37°C for 20 min to remove epithelial cells, and then minced and dissociated in RPMI containing collagenase (1 mg/ml collagenase II; Roche), DNase I (100 µg/ml; Sigma), dispase (0.05 U/ml; Worthington) and 10% FBS with constant stirring at 37°C for 45 min (SI) or 60 min (LI). Leukocytes were collected at the interface of a 40%/80% Percoll gradient (GE Healthcare). For transcription factor analysis, lamina propria mononuclear cells were first stained for surface markers before fixation and permeabilization, and then subjected to intracellular TF staining according to the manufacturer’s protocol (eBioscience Intracellular Fixation & Permeabilization buffer set from eBioscience).

### Sorting of intestinal ILCs

NK, ILC1, ILC2, and ILC3 (CCR6^+^ and CCR6^-^) were sorted from the small or large intestine lamina propria preparation using the surface marker panel described in **Figure S1**. A portion of sorted cells was stained intracellularly with ILC lineage transcription factors to confirm that the correct populations were isolated. Lin^-^Klrg1^hi^ cells were uniformly Gata3^hi^ and thus mature ILC2 (Hoyler et al., 2012). We have found Klrb1b to be a good marker of ILC1 and ILC3, which can then be further stratified based on expression of NK1.1^hi^ (ILC1) and NK1.1^lo^-NK1.1^neg^ (ILC3) (**Figure S1**). ILC1 are T-bet^hi^ while ILC3 are RORgt^+^ and can be either T-bet^-^ (CCR6^+^) or T-bet^+^ (CCR6^-^) (**Figure S1**).

### Cell staining for flow cytometry

For analysis of ILCs, live LP cells were stained with anti-NK1.1, anti-IL7ra, anti-Klrg1, anti-Klrb1b (clone 2D12, subclone from 2D9) as previously described (Aust et al., 2009), and anti-CCR6. Anti-CD11b, anti-CD11c, anti-CD14, anti-CD19, anti-B220, anti-TCRβ, anti-TCRγδ, and anti-CD3 were used as dump gate. For transcription factor staining, cells were stained for surface markers, followed by fixation and permeabilization before nuclear factor staining according to the manufacturer’s protocol (FOXP3 staining buffer set from eBioscience) with individually or in combination anti-RORgt, anti-T-bet, and anti-Gata3. 4′,6-diamidino-2-phenylindole (DAPI) or Live/dead fixable blue (ThermoFisher) was used to exclude dead cells. Flow cytometric analysis was performed on an LSRII (BD Biosciences) or an Aria (BD Biosciences). All data were re-analyzed using FlowJo (Tree Star). Commercially available antibodies used in flow cytometry experiments above are listed in **Table S1**.

### ATAC-seq

Sorted ILC populations were prepared according to (Buenrostro et al., 2013). Paired-end 50bp sequences were generated from samples on an Illumina HiSeq2500. Sequences were mapped to the murine genome (mm10) with bowtie2 (2.2.3), filtered based on mapping score (MAPQ > 30, Samtools (0.1.19)), and duplicates removed (Picard). For each sample individually, we ran Peakdeck (parameters –bin 75, -STEP 25, -back 10000, -npBack100000) and filtered peaks with P_raw_<1E-4. To enable quantitative comparison of accessibility across samples, we generated a reference set of accessible regions, taking the union (Bedtools) of peaks detected in individual samples. The reference set of ATAC-seq peaks contained 65,281 potential regulatory loci, ranging from 100 base pairs (bp) to 3750 bp (median length 375 bp). Reads per reference peak were counted with HTSeq-count. This pipeline resulted in ~14 million uniquely mapped reads per sample, ~42% of which mapped to the identified peaks. We used DESeq2 to normalize and identify differentially accessible peaks across conditions. Downstream analysis and data visualization were performed in MATLAB R2014B. For PCA analysis (Figure 1), ATAC-seq data was treated as in (Karwacz et al., 2017) to yield ~127k accessible regions, and robustly normalized DESeq2 counts were mean-centered and variance-normalized by the square root of standard deviation. ATAC-seq data were deposited in the GEO database under accession GSE116093.

### RNA-seq

For the initial RNA-seq measurements in wildtype ILCs, cells were sorted (FACSAria II, BD) directly into Trizol and snap-frozen, thawed for chloroform extraction and purified using the RNAeasy minElute kit (Qiagen). Samples were evaluated for RNA quality by sampling and analyzing one µl of library by Bioanalyzer (Agilent, Santa Clara, CA), using DNA high sensitivity chip. An accurate quantification of library concentration, was performed by NEBNext Library Quant Kit (New England BioLabs). Libraries were prepared using Nugen Ovation RNAseq System V2 and Nugen Ovation Ultralow Library System kits and sequenced on an Illumina HiSeq2500. Sequences were mapped to the murine genome (mm10) with STAR. HTSeq-count was used to count reads per gene. DESeq2 was used to normalize and identify differentially expressed transcripts among ILC lineages. Additional ILC RNA-seq data were downloaded from GEO: GSE85152, GSE77695, and processed similarly. We controlled for batch effects across the three datasets using COMBAT, yielding 62 samples in the final gene expression matrix. The combined samples clustered by ILC subsets (data not shown) and exhibited similar lineage-specific gene signatures for TFs, receptors and cytokines (**Figures S5-S7**). The full batch-corrected gene expression values for the combined studies is contained in **Table S3**.

For RNA-seq of *Bcl6* KO and control animals, ILC populations were sorted as described above. Briefly, cells were sorted into 100% FBS, and immediately spun at 600 g for 6 min. After removal of supernatant, the cell pellet was lysed with Lysis/Binding Buffer from mirVana miRNA Isolation Kit (Thermo Fisher, Grand Island, NY). Samples were evaluated for RNA quality as described above. The library for RNA-seq was prepared by using NEBNext Ultra II Directional RNA Library Prep kit (New England BioLabs, Ipswich, MA). To study differential gene expression, individually indexed and compatible libraries were proportionally pooled (~25 million reads per sample in general) for clustering in cBot system (Illumina, San Diego, CA). Libraries at the final concentration of 15 pM were clustered onto a single read (SR) flow cell v3 using Illumina TruSeq SR Cluster kit v3, and sequenced to 51 bp using TruSeq SBS kit v3 on Illumina HiSeq system. Sequences were mapped and gene expression analysis performed as described above.

RNA-seq data were deposited in the GEO database under accession GSE116093.

#### Core ILC subset gene sets

ILC subset gene sets were used to develop core networks (Figure 3) and for detection of lineage regulators (Figure 5). They were constructed from our gene expression dataset (20 samples). For a given lineage, ILC subset genes were included in the set if they were more highly expressed (log_2_(FC)>1, FDR=10%) in that subset relative to at least one other subset and not decreased (log_2_(FC)<-1, FDR=10%) relative to any other subset in the SI. This analysis yielded overlapping subset-defining gene sets ranging from ~1500-~2500 genes per subset. We also define sets of subset-suppressed genes, which, for a given subset, had to be less expressed in that subset relative to at least one other subset and not increased relative to any other subset in the SI. These core ILC gene signatures of up- and down-regulated genes are contained in **Table S3.**

### Transcriptional regulatory network inference

#### Motif Analysis and Generation of ΔATAC Prior Matrix

Peaks were associated with putative transcription-factor (TF) binding events and target genes to generate a “prior” network, *P* ∈ ℝ^|*genes*|×|*TFs*|^, of TF-gene interactions. We collected mouse motifs for 859 TFs from CisBP (Weirauch et al., 2014) and ENCODE (Kheradpour et Kellis, 2014), as described in (Miraldi et al., 2018). We scanned peaks for individual motif occurrences with FIMO (parameters –thresh .00001, –max-stored-scores 500000, and a first-order background model) and included motif occurrences with P_raw_<1E-4 in our analysis. To construct the ∆ATAC prior, core ILC subset ATAC-seq peak sets were defined based on differential accessibility, analogously, with the same parameters, to the core ILC subset gene sets above. For each subset and core TF, we began with all ATAC-seq peaks containing at least one motif occurrence (as the background set), we then tested whether core subset ATAC-seq peaks were enriched nearby core subset genes. Note that to perform the enrichment, genes were mapped to motif-containing peaks; gene sets were converted to “peak sets” and included the union of motif-containing peaks falling within +/-10kb of a gene body in the gene set. We used the hypergeometric CDF to estimate enrichment and applied a nominal significance cutoff of P_raw_<1E-4. TFs significantly enriched in ∆ATAC proximal to core gene sets were included as regulators of the genes in the set that contained a ∆ATAC-proximal TF-motif-containing peak. To detect repressive interactions, we repeated the procedure above looking for enrichment of core ATAC-seq peaks nearby subset-suppressed genes. This procedure was also repeated for gene sets derived from CCR6^-^ ILC3, CCR6^+^ ILC3 and ILC2 samples from the LI. Finally, we also derived LI- and SI-dependent TF-gene interactions, similarly testing for enrichment between ∆ATAC and Dgene sets based on LI versus SI accessibility / expression in the three cell types above. The resulting “∆ATAC” prior matrix contained 42,850 TF-gene interactions for 4226 genes and 129 TFs (**Table S4**).

#### Inference framework

We built gene expression models according to (Miraldi et al., 2018), using a modified LASSO-StARS framework to solve for TF-gene interaction terms {bik}:

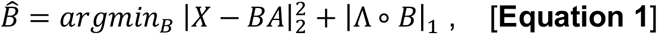

where, *X* ∈ ℝ^|*genes*|×|*samples*|^ is the expression matrix for genes in the prior, *A* ∈ ℝ^|*TFs*|×|*samples*|^ contains TF activities, B ∈ ℝ^|*genes*|×|*TFs*|^ is the matrix of nonnegative penalties, and ∘ represents a Hadamard (entry-wise matrix) product (Gustafsson et al., 2015; Studham et al., 2014). Representing the LASSO penalty as a Hadamard product involving a matrix of penalty terms, as opposed to a single penalty term, enabled us to incorporate prior information into the model-building procedure. Specifically, a smaller penalty Λ_*ik*_ is used if there is evidence for the TF-gene interaction in the prior matrix. This procedure encourages selection of interactions supported by the prior (e.g., containing ATAC-seq evidence), if there is also support in the gene expression data.

For this study, our target gene expression matrix contained the 6418 “core” ILC genes (defined above) for the 62 samples. We considered 445 potential TF regulators (intersection of “core” ILC genes with our curated list of mouse TFs (Miraldi et al., 2018)). Entries of the Λ matrices were limited to two values: the nonnegative value *λ*, for TF-gene interactions without evidence in the prior, and .25 × *λ*, for TF-gene interactions with support in the prior. Note that 25 × *λ* weights interactions in the prior twice as strongly as in our previous Th17 TRN (Miraldi et al., 2018), because, in this context, we have fewer gene expression samples (62 versus 254) and a presumed higher confidence prior (due to ∆ATAC method above).

As described previously, we used the stability-selection method StARS (Liu et al., 2010) with 50 subsamples of size. 63 × |*samples*| and an instability cutoff = .05 to solve for the TF-gene interactions B. We then ranked TF-gene interactions based on stability as described in (Miraldi et al., 2018) and used out-of-sample prediction (described below) to determine how many interactions to include in the final TRN. To estimate the TF activity matrix *A*, we used two methods: (1) TF gene expression and (2) prior knowledge of TF target gene expression (Arrieta-Ortiz et al., 2015). For (2), we used the following relationship to solve for TF activities: *X = PA* [**Equation 2**], where *P* ∈ ℝ^|*genes*|×|*|TFs*|^ is the (ΔATAC) prior. Based on previous results (Miraldi et al., 2018), we generated separate TRNs using each of the TF activity methods and rank-combined TRNs using the maximum. Based on gene-expression prediction (described below, **Figure S4D,E**), our ILC TRN size was set to an average of 10 TFs per gene (total 63,832 TF-gene interactions). At this model size, 25,113 TRNs interactions (39%) had ∆ATAC prior support, and the TRN contained 59% of the 42,850 prior interactions. Thus, many ΔATAC are filtered out (41% of the prior) because they are not supported by the gene expression data and new interactions (61% of the final TRN) are learned from gene expression patterns alone.

#### Gene Expression Prediction

TRN model quality was computationally assessed by out-of-sample gene expression prediction. To ensure robustness of results, four separate out-of-sample gene expression prediction tasks were designed, where, for each independently, ILC TRN models were built in the absence of (1) all ILC1 (5 samples) from (Gury-BenAri et al., 2016), (2) all ILC2 (5 samples) from (Gury-BenAri et al., 2016), (3) all ILC3 (6 samples) from (Gury-BenAri et al., 2016), and (4) all eight large intestine (LI) samples. For each out-of-sample prediction challenge, models were built using prior-based TFA (TFA=P^+^A) and TF mRNA (see “*Inference Framework*”). Because the LI gene expression data had been used to construct the ∆ATAC prior, TF-gene interactions based on gene sets derived from the LI samples were omitted and the reduced prior was used to train the model.

Training TFA matrices were mean-centered and variance-normalized according to the training-set means *ā*_*train*_ ∈ ℝ^|*TFs*|^ and standard deviations 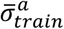 ∈ ℝ^|*TFs*|^. Target gene expression vectors were mean-centered according to the training-set mean *x̅*_*train*_ ∈ ℝ^|*genes*|^. Predictive performance was quantified with 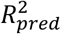 as a function of mean model size (TFs / gene) (**Figure S4D,E**). In brief, for each model-size cutoff, we regressed the vector of normalized training gene expression data onto the reduced set of normalized training TFA estimates to arrive at a set of multivariate linear coefficients *B*_*train*_ ∈ ℝ^|*genes*|×|*|TFs*|^. 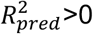 is defined as:

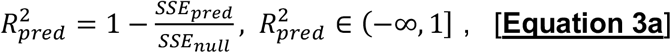

where

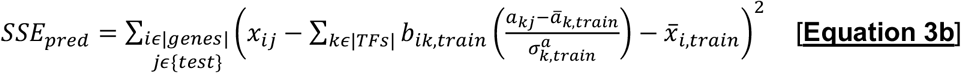

and

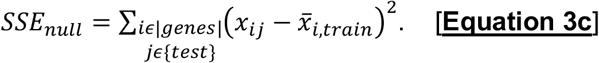

Any 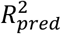>0 indicates that the numerator model has predictive benefit over the null model (simply using mean gene expression from the observed training data). For all leave-out gene expression prediction tasks, median 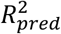 increased dramatically from model-size two to five TFs / gene and began to plateau at 10 TFs / gene, suggesting 10 TFs / gene as model-size cutoff (**Figure S4D,E**). We note that ILC TRN analyses (Figures 3-5, 7) were tested for robustness to variation in this cutoff (e.g., **Figure S7A**).

### Network Visualization

Networks were visualized using a newly designed interactive interface, based on iPython and packages: igraph, numpy and scipy (https://github.com/simonsfoundation/jp_gene_viz). ILC TRNs are available from the following link: https://github.com/flatironinstitute/ILCnetworks (see **<u>Supplemental Text 1</u>** for detailed instructions).

### TF-TF Module Analysis

TF-TF modules were constructed as described in (Miraldi et al., 2018). In brief, for positive and negative edges separately, we calculated a background-normalized overlap score z_*ij*_ between TF *i* and TF *j* as:

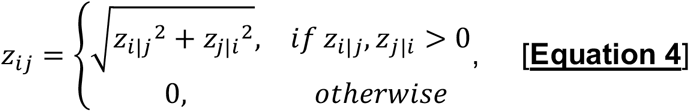

where z_*i*|*j*_ is the z-score of the overlap between TF *i* and TF *j*, using the mean and standard deviation associated with the overlaps of TF *j* to calculate the z-score. We filtered the normalized overlap matrix so that it contained only TFs with at least one significant overlap (FDR = 10%). We then converted the similarity matrix of normalized overlaps to a distance matrix for hierarchical clustering using Ward distance. To arrive at a final number of clusters, we calculated the mean silhouette score for solutions over a range of total clusters and selected the solution that maximized silhouette score. For positive interactions, this analysis lead to 57 clusters, and 12 clusters for negative interactions. We estimated the significance of individual clusters based on cluster size and overlap between TFs in the cluster (Miraldi et al., 2018). We only report positive TF-TF clusters, because, as observed previously (Miraldi et al., 2018), positive TF-TF clusters were orders of magnitude more significant than TF-TF clusters based on repressive interactions.

## Supplementary Table Legends

**Table S1.** Commercially available antibodies used in flow cytometry experiments.

**Table S2.** Batch-corrected gene expression data combining samples from this study, Gury-BenAri et al., and Shih et al.

**Table S3.** Up-regulated and down-regulated ILC subset gene signatures (see **Methods**).

**Table S4**. ∆ATAC prior matrix.

**Table S5.** The ILC TRN.

